# Heavy metal resistance promotes higher biomass formation in multidrug-resistant wastewater *Escherichia coli* isolates from Finland

**DOI:** 10.64898/2026.03.16.712119

**Authors:** Ahmad Ibrahim Al-Mustapha, Riikka Laukkanen-Ninios, Kirsi-Maarit Lehto, Päivi Tammela, Annamari Heikinheimo, Cristina D. Cruz

## Abstract

Antimicrobial resistance alone is a serious threat to global public health, but even more so when multidrug-resistant bacteria harbour heavy metal resistance genes, as these can drive co-selection of antibiotics and produce biofilms. The presence of such genes, combined with the bacteria’s ability to form biofilms, is strongly linked to treatment failure, persistent infections, and reduced therapeutic options. Here, we used the resazurin-crystal violet combination assay to screen a representative cohort of whole-genome sequenced *Escherichia coli* isolates (n=20) obtained from wastewater surveillance. The specific biofilm formation (SBF) index was used to grade the intensity of biofilm formation as strong, moderate, weak, and non-biofilm producers. Correlation analysis was used to test the association between the intensity of biofilm formation and genotypic features. The SBF index revealed that most of the wastewater *E. coli* isolates (n=13/20) were weak/non-biofilm producers, four isolates produced moderate biofilms, and three isolates (ST1434: A-O18ab:H55; ST401:A-H25; and ST399:A-O19:H12) produced strong biofilms. The diversity of virulence factors was similar in most of the isolates, except for the three isolates, which had fewer abundant virulence factors. The correlation analysis showed that there was no association between the expression of virulence genes and the formation of strong biofilms by the isolates (p > 0.05). Drug resistance profile was not correlated with higher biofilm production (*p* > 0.05), as 68.8% (n=11/16) of multi-drug resistant (MDR) and 50% (n=2/4) of non-MDR isolates had weak or no biofilm formation. Similarly, the SBF index was not associated with the number of plasmids in each of the *E. coli* genomes (*p* = 0.334). However, there was a positive association between the presence of two or more heavy metal resistance genes (HMRGs) and the strong biofilm formation in our isolates (*p* = 0.002). Our findings revealed the low occurrence of strong biofilm producers among wastewater *E. coli* isolates. Further studies are needed to evaluate the impact of the presence of HMRGs and their direct or indirect contribution to enhancing biofilm production and persistence in environmental reservoirs.

## Introduction

Resistance mechanisms such as the acquisition of antimicrobial resistance genes, disinfectant resistance, and biofilm formation are some of the key mechanisms of bacterial survival.^1-3^ Biofilms are structured microbial communities encased in a self-produced extracellular polymeric matrix, which adhere to biotic or abiotic surfaces, including medical devices and biomaterials.^4,5^ These complex aggregates are a significant concern in clinical and industrial settings due to their inherent resistance to antimicrobial agents and host immune responses.^6,7^ *Escherichia coli* (pathogenic or commensal) are gram-negative, biofilm-forming pathogens associated with persistent infections, medical device contamination, and foodborne illnesses.^6^

The traditional techniques for biofilm assessment by crystal violet (CV) staining measure adherent biomass by binding to extracellular polysaccharides and cellular components. While widely used, CV staining alone does not provide information on metabolic activity within the biofilm. Conversely, the resazurin assay (also known as the Alamar Blue assay) evaluates metabolic activity by detecting the reduction of resazurin to fluorescent resorufin in viable cells.^9^ The resazurin-CV combination assay offers a more comprehensive phenotypic evaluation of biofilm formation by quantifying both metabolic activity and biomass, respectively.^4,8-10^

The emergence and spread of multidrug-resistant (MDR) and high biofilm-producing bacterial pathogens pose a threat to the control of bacterial infections in humans and animals. ^4,10^ The study of antimicrobial-resistant pathogens has significantly advanced with the availability of genomic technologies like whole genome sequencing (WGS). We have earlier published a WGS-based genotypic profile of a cohort of *E. coli* isolates that were obtained from the influents of ten wastewater treatment plants in Finland.^11^ Our findings from that study revealed the spread of MDR AmpC/ESBL-producing *E. coli* isolates in the Finnish population, and some of the isolates were genotypically similar to human surveillance isolates also collected from Finland.

Studies have also shown that the wastewater pipeline could impact the fate, genotype, and resistance profiles of isolates obtained from wastewater.^12^ Environmental bacteria have demonstrated a remarkable capacity to adapt to toxic conditions, including environments contaminated with heavy metals. This adaptation often involves the development of resistance mechanisms to heavy metal ions, such as efflux pumps, enzymatic detoxification, and sequestration processes.^2^ The situation becomes increasingly concerning when these bacteria simultaneously acquire antibiotic resistance, a phenomenon widely reported in both clinical and environmental isolates.^3^ The association between biofilm formation and heavy metal resistance has been previously reported for both gram-positive and gram-negative bacterial species isolated from food products.^1^ Hence, our objective for this paper was to evaluate the ability of biofilm formation among a subset of environmental *E. coli* isolates and assess which of the genotypic features (antibiotic resistance profiles, virulence genes, plasmids) obtainable from WGS in-silico analysis could predict the biofilm formation intensity. The findings of this study are expected to enhance the understanding of the biofilm-forming status of wastewater *E. coli* strains and their public health and environmental implications. Our preliminary data suggest a possible association between biofilm formations and heavy metal resistance, which has been previously shown for gram-negative species isolated from dairy and non-dairy products.^1^

## Materials and Methods

### *Escherichia coli* strains and growth conditions

A total of 20 representative previously whole-genome sequenced *E. coli* isolates were included in this study. The isolates were obtained from wastewater surveillance for MDR bacteria as part of the WastPan project in Finland.^13^ During the study, 75 *E. coli* isolates were obtained from influents of ten wastewater treatment plants (WWTPs) across Finland between February 2021 and January 2022 and were stored at -80°C.^11^

For this study, we purposively selected the isolates to encompass most of the Multilocus Sequence Types, city of isolation, and antibiotic resistance profiles (resistant and sensitive strains). Each of the *E. coli* isolates was subcultured on Lysogenic broth (Oxoid, Basingstoke, UK) plates and cultured at 37°C overnight (18-24 h) and stored at 2 - 4°C as the monthly working culture stock. A weekly working culture was subsequently used for making the bacterial suspension by suspending distinct bacterial colonies in 5 mL of 0.9% sterile saline solution. The turbidity of the suspension was measured using a densitometer (DEN-1, BioSan, Warren, Michigan, USA), and the bacterial inoculum, adjusted to 1×10^6^ colony-forming units (CFU)/mL, was added (200 μL) into wells of a 96-microtitre plate (Nunc, Roskilde, Denmark). LB broth (Oxoid, Basingstoke, UK) was included in the same plate and used as a negative control. The microtiter plate was then incubated for 24 h at 37 °C without shaking to ensure the formation of mature biofilm. *E. coli* ATCC 25922 (ST10:B2-O6:H1), a known weak biofilm producer, was used as a control strain for the combination assay.

### Resazurin and Crystal Violet Combination Assay

The combination assay was conducted in 96-well microtiter plates. Each isolate was screened in six replicates, and three independent assays were conducted. Bacterial growth is measured in the Multiskan GO plate reader (Thermo Fisher Scientific, Vantaa, Finland) at Time 0 and after incubation (18-24 hours) prior to the resazurin assay. The resazurin assay was used for the assessment of metabolically active cells. Resazurin (7-hydroxy-3H-phenoxazin-3-one 10-oxide) is a blue dye, itself weakly fluorescent until it is irreversibly reduced to pink-colored and highly fluorescent resorufin. To start the resazurin assay, the overnight culture media was removed, and 200 µL of 1× phosphate-buffered saline (PBS) was added to wells to wash off and remove non-adherent, planktonic cells. PBS was removed, and 200 µL of freshly prepared resazurin solution (Sigma-Aldrich, diluted to 4 µg/mL in sterile 1× PBS) was added to each well, and the plates were immediately covered with aluminum foil. The plates were incubated at 37°C for 180 min. The fluorescence (λex = _570nm_, λem = _615nm_) was measured using a multimode microplate reader Varioskan LUX (Thermo Fisher Scientific, Vantaa, Finland).

As opposed to the resazurin assay, the CV assay stains not only cells but essentially any material adhering to the surface of the plate (e.g., matrix components). To proceed with the combination assay, the resazurin solution was carefully removed from the wells using a multichannel pipette without disturbing the biofilm. The biofilms were then fixed by adding 200 µL of 99.5% ethanol and incubated at room temperature (RT) for 5 min. The ethanol was removed, and the plates were allowed to air-dry for 10 min. Thereafter, 200 µL of 0.01% crystal violet aqueous solution (Sigma-Aldrich) was added to the wells, and the plates were incubated for 30 min at RT. The stain was then removed, and the plates were washed three times using 210 µL of sterile water, after which the plates were air-dried (mostly for 30-45 minutes, depending on the volume of water left in each well during the wash stage). A final elution of the biofilms was done by adding 200 µL of 94% ethanol to each well to dissolve the CV stain, and the plates were incubated for 1 h at RT. Finally, the absorbance at OD_595nm_ was measured using a Multiskan GO plate reader (Thermo Fisher Scientific, Vantaa, Finland). Three independent experiments were performed with six replicates each.

### Data Analysis

All statistical analysis was done in SPSS *v*.*28*. The quantitative variables were presented as mean and standard deviation. The specific biofilm formation index (SBF) was calculated based on the recommended cut-off by Naves et al.^14^ using the formula: SBF = AB-CW/G. AB is the OD_595nm_ of stained attached bacteria, CW is the OD_595nm_ of stained control wells containing bacteria-free medium only, and G is the OD_595nm_ of cell growth in suspended culture. The SBF was thereafter graded as: strong biofilm producers with an SBF ≥1.10; moderate producers with an SBF of 0.7-1.09; weak producers with SBF of 0.35-0.69; and no biofilm producers with SBF of ≤0.34.^14^ Correlation analysis between biofilm formation, and genotypic features (virulence factors, number of antibiotic/stress resistance genes, and number of plasmids) were computed using Pearson’s r coefficient with a significance of *p* values < 0.05. The number of plasmids and ARGs was previously described by Al-Mustapha et al.^11^ In this study, the virulence genes were identified using the *E. coli* Virulence Factor Database (VFDB) tool in Ridom Seqsphere+ *v*. 10.5.

## Results

### Metabolic activity and biomass production

The resazurin assay revealed that all the cultured bacteria were metabolically active, with a range of 90.21 and 896.2 relative fluorescence units (RFU). The assay revealed significant variations in the proportion of metabolically active cells within the biofilm in each of the samples (six replicates each), with a mean of 551 ± 159.6 (Figure 1). The lowest metabolically active isolate was AE08 (ST10:D-H9), which was a slow grower, while AE54 (ST399:A-O19:H12) was the isolate with the most metabolically active cells (Figure 1).

**Figure 1.**
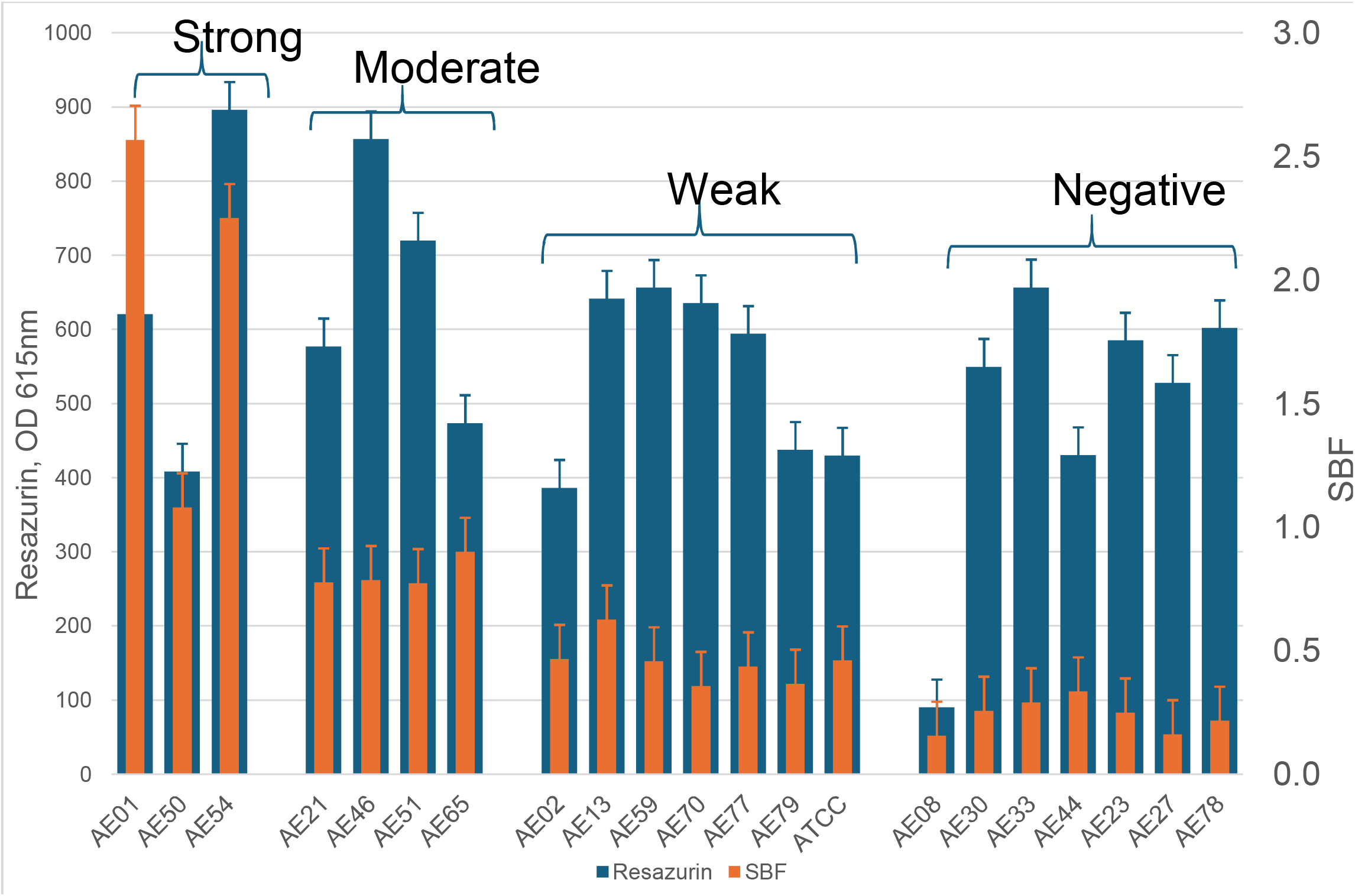
Resazurin-crystal violet combination assay of 20 *Escherichia coli* samples in Finland. SBF-Specific biofilm formation index. Data represent the mean ± standard deviation from three independent experiments (n=6). For detailed genomic characteristics of the isolates, see **Table 1**.

The CV assay revealed that 65% (n=13/20) of the wastewater *E. coli* isolates were weak or non-biofilm producers. Amongst these, six of them were graded as non-biofilm producers (i.e., SBF ≤0.35, Figure 1). Four isolates (AE21-ST2619:D-O1:H7, AE46-ST69:B2-H4, AE51-ST405:A-O102:H6, and AE65-ST38:F-O86:H18) produced moderate biofilms with an SBF range of 0.78 – 0.90. Three isolates belonging to phylogroup A (AE01-ST1434:O18ab:H55, AE50-ST401:H25, and AE54-ST399:O19:H12) produced strong biofilms (Figure 1; Table 1).

**Table 1.**
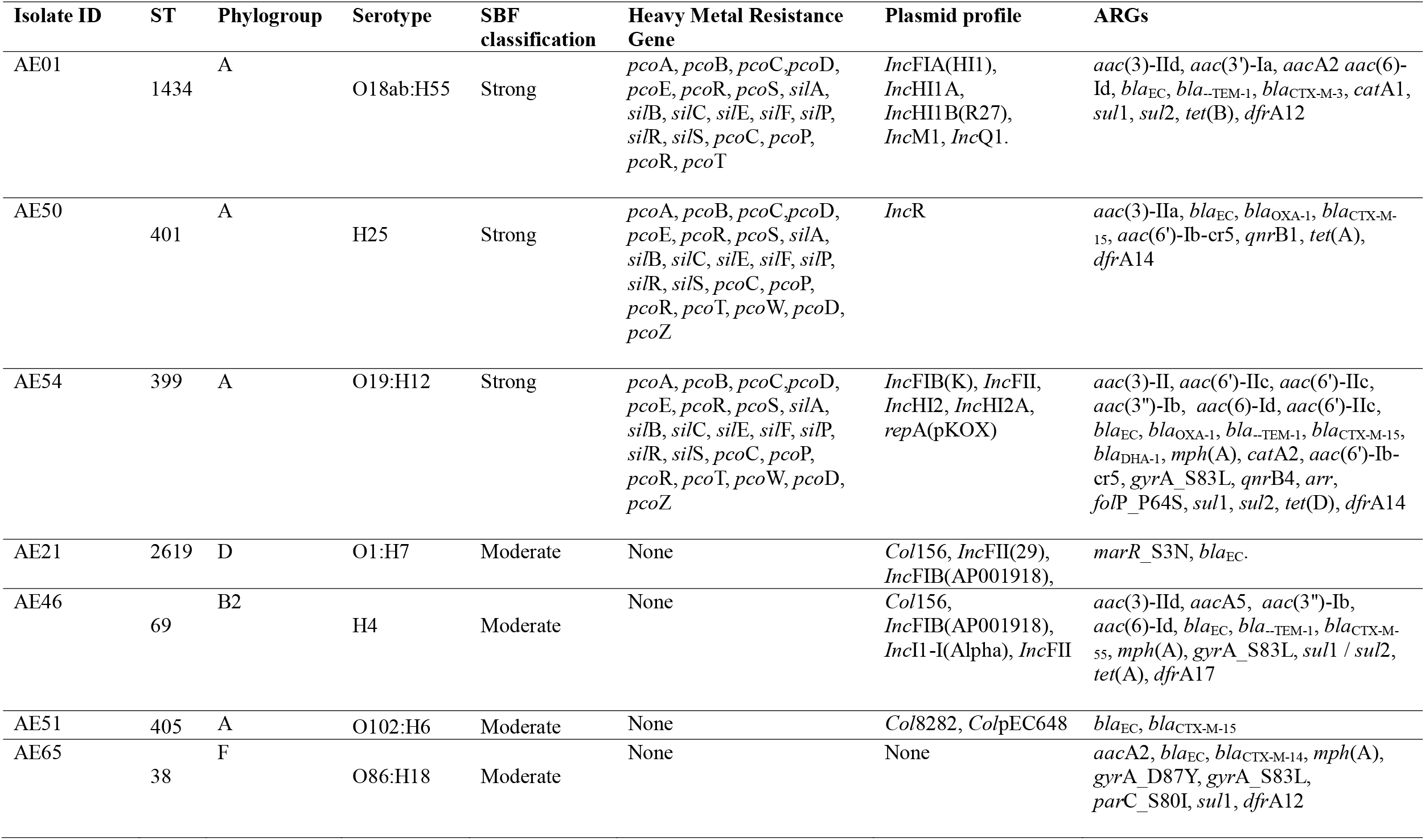
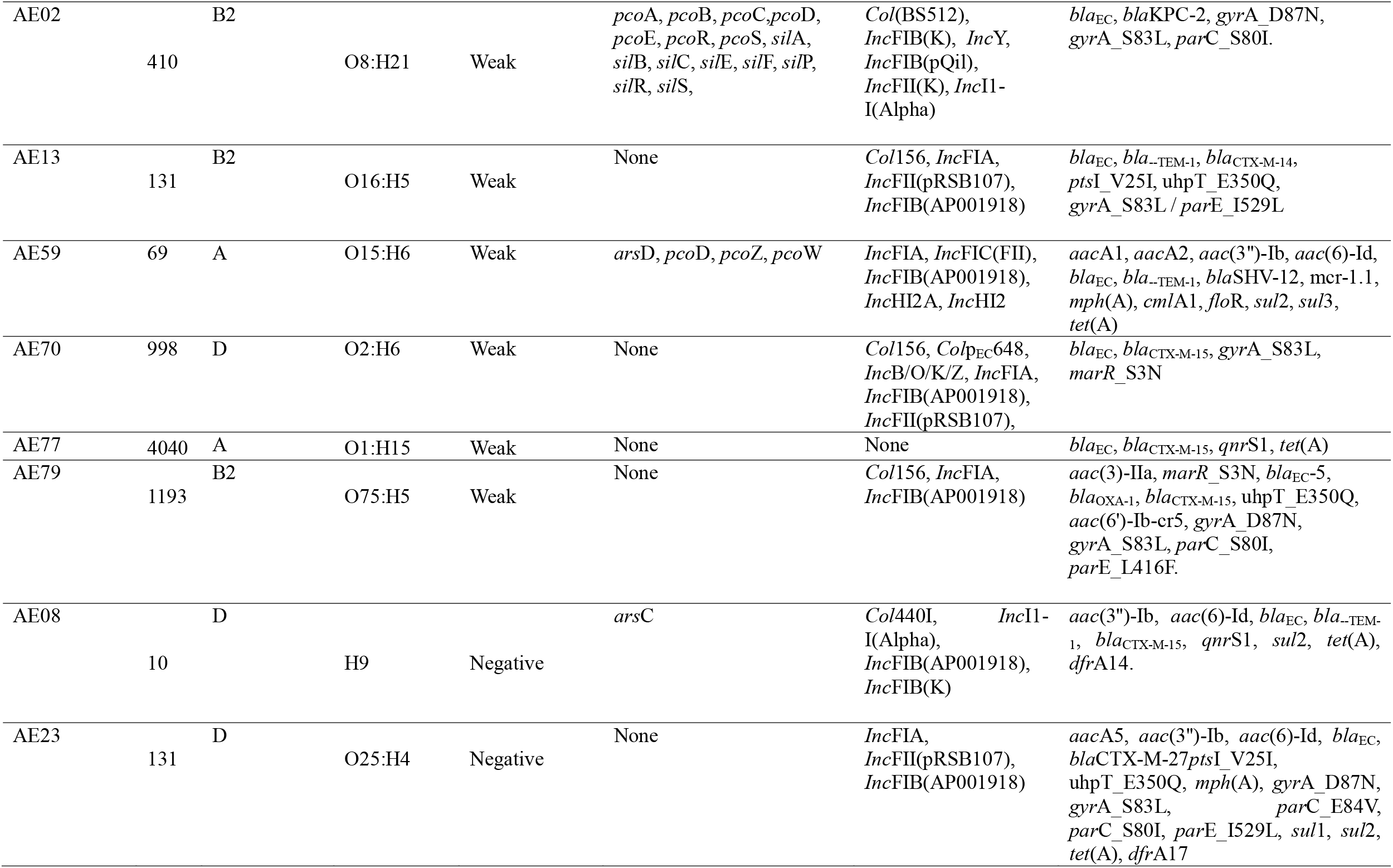
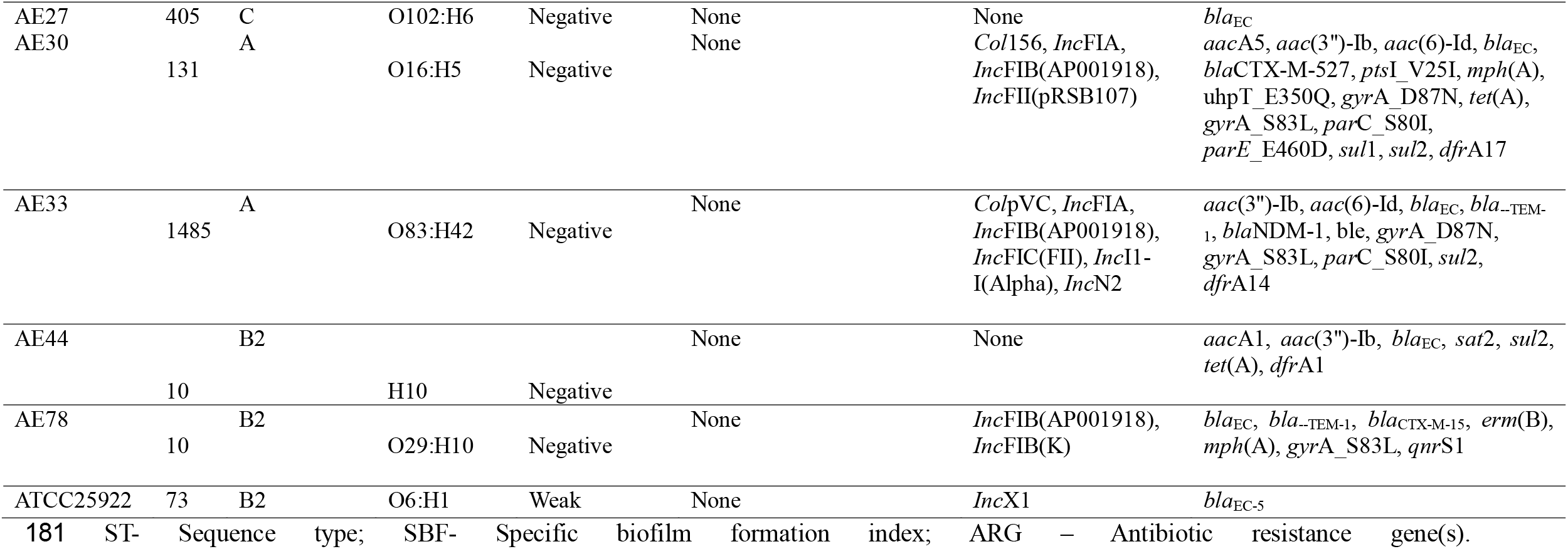
Genotypic and biofilm-forming profiles of wastewater of *Escherichia coli* isolates (n=20).

### Genotypic profiles and their association with biofilm formation

Analysis of the WGS revealed that the 20 isolates harboured several virulence factors that could be grossly categorized into those associated with the type III secretion system, such as *esp*X4, *esp*X5, *esp*Y3, *esp*Y4; bacterial adhesion and initial attachment (such as the *fim*ABCDEFGHI operon, the *pap*BCDGIX operon, fdeC, *yag*KVWXYZ/*ecp*ABCDER operon); and iron uptake (*iuc*ABC, *chu*ASTUVWXY, *ent*BCDEF, and *fep*ABCDG). In addition, some isolates harboured the *asl*A, *kps* genes (*kps*DMT), and *omp*A, which are crucial to the extracellular matrix and biofilm stability, and several toxins (*sat, vat*, hlyABCD, cnf-1, *set*1AB, and *tcp*C). The diversity of virulence factors was similar in most of the isolates, except for the three strong biofilm producers (AE01, AE50, and AE54), which had fewer abundant virulence factors. These isolates only have two groups of genes related to bacterial adhesion (*esp*X and *fim*), two genes related to iron uptake (*ent* and *fep*) and a single gene related to the formation of the extracellular matrix (*omp*A) (Figure 2). These isolates also had at least three HMRGs (mercury, copper, and silver), and several plasmid replicons (AE01 and AE54). There were no correlations between the expression of most virulence genes and the formation of strong biofilms by the isolates (p > 0.05).

**Figure 2.**
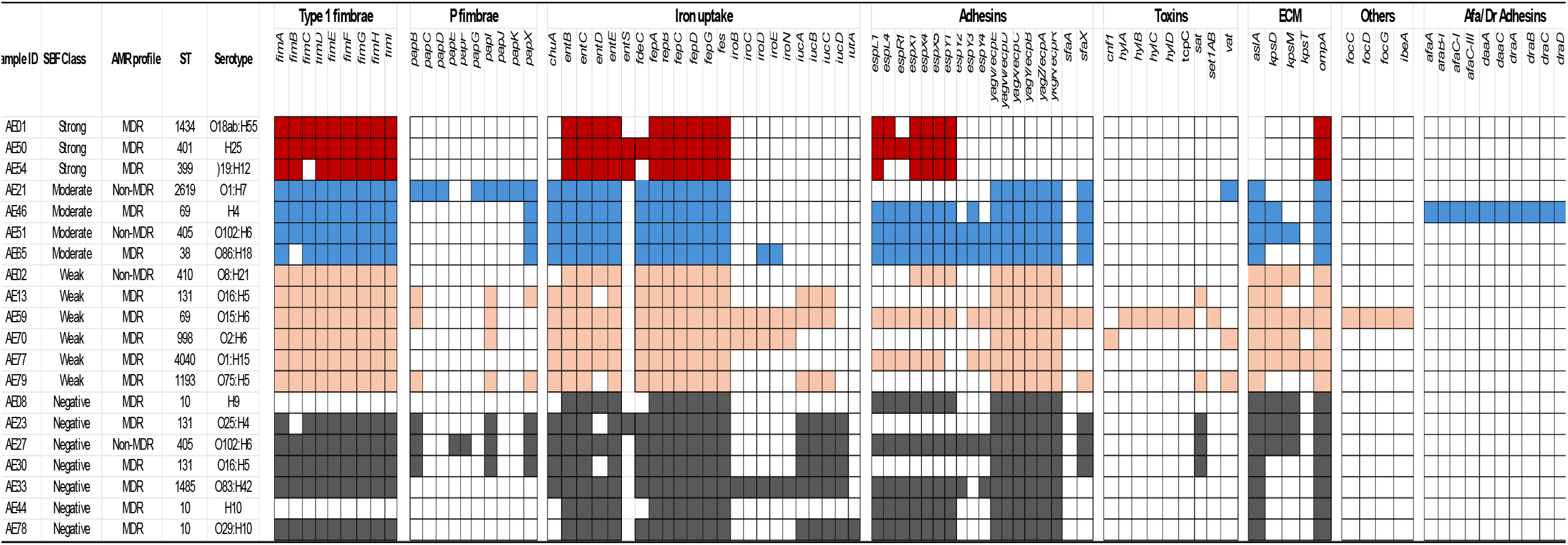
SBF – Specific biofilm formation, AMR profile, and genomic traits associated with wastewater *E. coli* collection (n=20) studied. The virulence factors were detected by WGS using the VFDB database. Based on the SBF index classification, strains were colour-coded as strong biofilm (red), moderate biofilm producers (blue), weak biofilm producers (light orange), and non-biofilm forming isolates (grey). ECM-genes associated with the formation of the extracellular matrix. Virulence determinants: *afa*A: transcriptional regulator; *afa*B: periplasmic chaperone; *afa*C: Outer membrane usher protein; *afa*D: afimbrial adhesin; *afa*E: adhesin protein; *chu*A: outer membrane hemin receptor; cnf1: cytotoxic necrotizing factor; hylABCF: hemolysin F; ibeA: invasin of brain endothelial cells; *iro*B; *iro*C, *iro*D, *iro*E, *iro*N: *ent*erobactin siderophore receptor protein; *iuc*C: aerobactin synthetase; iutA: ferric aerobactin receptor; *kps*E: capsule polysaccharide export inner-membrane protein; *kps*M: polysialic acid transport protein group 2 capsule; *omp*T: outer membrane protease *pap*A: Major pilin protein; *pap*C: outer membrane usher P fimbriae; pic; *sat*: secreted autotransporter toxin; senB: plasmid-encoded *ent*erotoxin; terC: tellurium ion resistance protein; *vat*: vacuolating autotransporter toxin; *yag*V/*ecp*: fimbrial protein.

Of the 20 isolates, four isolates (AE02, AE21, AE27, and AE51) had two or no known antibiotic resistance genes (ARGs) and were non-MDR. All the other 16 isolates harboured ARGs belonging to three or more antibiotic classes and were grouped as MDR. The MDR status was not correlated with a higher biofilm production (*p* > 0.05), as 68.8% (n=11/16) and 50% (n=2/4) of MDR and non-MDR isolates, respectively, exhibited weak or no biofilm formation. Likewise, the SBF index was not associated with the number of plasmids in each of the *E. coli* genomes (r^2^ =0.242; *p* = 0.334). Several stress resistance genes were also detected in the isolates, with the resistance to heavy metals being the most notable. Pearson’s correlation revealed a strong positive association between the presence of two or more HMRGs and the strong biofilm formation in our isolates (r^2^ =0.68; *p* = 0.002).

## Discussion

Biofilm formation is an important pathogenic determinant and a universal bacterial survival strategy,^1,14^ thereby facilitating bacterial persistence. Our study revealed that three of the isolates studied produced strong biofilm, and four isolates produced moderate biofilm. We therefore agree with studies that reported low biofilm formation amongst environmental *E. coli* isolates and great variations in the ability of *E. coli* to form biofilms in vitro.^1,5,15^

The resazurin-CV combination assay is a very easy and reproducible method, ideal for higher throughput phenotypic screening of isolates.^4^ From the literature, most biofilm studies have been carried out on clinical isolates or laboratory strains. Our findings are unique as they present evidence of the critical role of wastewater as a reservoir for *Escherichia coli* strains harboring HMRGs and exhibiting biofilm-forming capabilities. The observed positive correlation between the presence of multiple HMRGs not only enhances bacterial persistence in the environment but also facilitates the horizontal transfer of ARGs, thereby increasing public health risks. Our findings, therefore, complement existing literature on biofilm formation, especially as the fate of wastewater isolates could vary considerably based on the variations in population carriage of the pathogen, the fate of the pathogen in the wastewater pipeline, the isolation technique, and its stress adaptation features.

The seven *E. coli* strains that produced strong or moderate biofilms were typed as O1, O18, O86, O102, and an untyped O serogroup. Our results agree with the report of Naves et al.^15^ with respect to serogroups O1 and O18 as being strong biofilm producers. Other authors reported that the members of serogroups O2, O4, O22 and O25 produce thicker biofilms on polystyrene plates than those of serogroups O1, O6, O16, O18, and O75.^16^

Several studies have reported a high correlation between the MDR status and biofilm formation among clinical *Enterobacteriaceae* isolates.^17,18^ A study reported that there was an association between resistance to expanded spectrum cephalosporin, cell surface proteins and biofilm formation when they transformed a *Salmonella* plasmid into an *E. coli* DH10B.^19^Another study reported that the analysis of biofilm formation among 137 clinical *K. pneumoniae* strains revealed that 85.0% of biofilm-positive strains could produce extended-spectrum ™-lactamases compared to biofilm negative strains with a rate of 11.7%.^20^ Other studies reported that extensively drug-resistant *K. pneumoniae* isolates also showed a greater ability to form a biofilm (91.1%) when compared to MDR and sensitive strains. ^21,22^ They found that the carbapenem resistance phenotype significantly correlated with the biofilm formation ability of *K. pneumoniae*, with 99.9% of carbapenem-resistant isolates forming moderate and strong biofilms. Our findings revealed that the two carbapenemase-producing isolates (AE02 and AE33) were weak or non-biofilm producers. Similar weak biofilm production among carbapenem-resistant isolates was previously reported.^23^ In general, our findings revealed that there was no correlation between the MDR status and biofilm formation among wastewater *E. coli* strains. Our findings were also supported by the reports of Zheng et al.^24^ and Sabenca et al.,^25^ who reported no significant association between biofilm formation and resistance to the examined antibiotics. The differences might be due to the differences between clinical and wastewater isolates, as well as geographical differences in sampling sites. However, more studies are needed to fully explore the association between AMR profiles and biofilm formation in *E. coli*.^26^

In general, genes of operons type 1 fimbrae *fimABCDEFGHI*, enterobactin *entE*BD, iron transporter *fep*ABCDG, and *fes*, as well as the *omp*A gene associated with extracellular matrix formation, were conserved in all three strong biofilm-producing genomes (Table 2). There were variations in the expression of all other genes, such as the six variants of the *E. coli* common pilus genes (*yag*V/*ecp*E, *yag*W/*ecp*D, *yag*X/*ecp*C, *yag*Y/*ecp*B, *yag*Z/*ecp*A, ykgK/*ecp*R), which were detected in all isolates except the strong biofilm producers (AE01, AE50, and AE54). There were no correlations between the expression of all other virulence genes and the formation of strong biofilms by the isolates.^27^ Markedly, while studies have reported an association between a higher potential to form biofilm and some urovirulence genes, none of the previously reported genes had a strong correlation with strong or moderate biofilm production among wastewater isolates. Our results were therefore not in line with the reports that *papC*, different *papG* alleles, *sfa/focDE, focG, hlyA, iroN*, and *cnf1* were more prevalent in the strong biofilm producers. The only isolate that harboured all of these virulence genes (except *pap*C) was a weak biofilm producer (AE70). ^15,16,27^

It was previously reported that strains of *E. coli* that express type 1 fimbriae were significantly more efficient in forming a biofilm.^28^ The type 1 fimbriae (*fim*A) were reported to play an important role in the initial attachment to abiotic surfaces during biofilm formation.^28^ However, our findings and those of Cornejova et al. ^27^ found *fim*A in all *E. coli* strains irrespective of their biofilm-forming abilities. Also, our findings were not in agreement with this assertion that haemolysin was associated with strong biofilm formation, as the only isolate (AE70) that had haemolysin genes (*hyl*A, hylB, hylC, hylD), which also harboured other toxins (*cnf*1, *set*1A, *set*1B, *tcp*C, and *vat*), was a weak biofilm producer with an SBF index of 0.36. The prevalence of the siderophore-related genes (*iroBCDEN, iucABCD/iutA*) was variable, but no significant differences were observed between strong, moderate, weak, and non-biofilm producers. Similarly, protectin/invasion-encoding genes such as *asl*A and *kpsDMT* were also widely distributed among moderate, weak, and non-biofilm-forming isolates. The *afa*ABCD (and their allelic variants), *daa*ACDF, and *dra*ABCDP adhesins were only detected in one moderate biofilm producer (AE46).

The increasing prevalence of heavy metal contamination in environmental and clinical isolates has exerted selective pressure on bacteria, leading to the emergence of some heavy metal-resistant strains.^30^ The three isolates that formed strong biofilms harboured three HMRGs: silver, mercury, and copper. This finding is supported by the report of Harrison et al., ^31^ which suggested a potential link between heavy metal resistance and enhanced biofilm formation. Also, exposure to sublethal concentrations of heavy metals has been shown to upregulate biofilm-related genes, increase exopolysaccharide production, and alter biofilm architecture. ^32^ HMRGs have also been associated with increased adhesion and biofilm stability.^33^ These observations raise the question: Does heavy metal resistance inherently promote increased biofilm formation, or is it a co-adaptive response to environmental stress? It is essential to explore potential molecular mechanisms, including the role of two-component regulatory systems, quorum sensing, and exopolysaccharide modulation in strong biofilm-producing and heavy metal-resistant strains such as the AE01 (ST1434: A-O18ab:H55). Our study has a few limitations. First, the sample size was small. Secondly, we did not determine phenotypic resistance to heavy metals.

In conclusion, we reported the low occurrence of strong biofilm producers among a cohort of 20 wastewater *E. coli* isolates. Our findings revealed a strong positive association between the presence of two or more HMRGs and strong biofilm formation in our isolates. Biofilms in wastewater systems serve as reservoirs for MDR pathogens, complicating treatment strategies and increasing the potential for outbreaks. Moreover potential link with HMRGs could increase co-selection of AMRs. More studies are needed to evaluate the impact of the presence of individual HMRG on biofilm production.

## Declarations

### Funding

This study utilized isolates obtained as part of the WastPan consortium project, funded by the Research Council of Finland (formerly the Academy of Finland) (grant number: 1339417), the Finnish Government, the Ministry of Agriculture and Forestry Finland, the Ministry of the Environment Finland, the Finnish Water Utilities Development Fund, and the National Emergency Supply Agency Finland. Funding from the Research Council of Finland (grant number: 362584) was used for the biofilm assays. The Helsinki University Library covered the open-access publication fee.

### CRediT authorship contribution statement

AH and AIA conceptualized the study. AIA and CDC conducted bacterial subculture and combination assay. AIA drafted the manuscript. AT, RLN, KML, AL, SO, and PT supervised the benchwork. All authors reviewed and approved the final manuscript.

### Declaration of competing interest

The authors declare that the research was conducted in the absence of any commercial or financial relationships that could be construed as a potential conflict of interest. All claims expressed in this article are solely those of the authors and do not necessarily represent those of their affiliated organizations or those of the publisher, the editors, or the reviewers.

### Data availability

No new data were generated.

### Ethics approval

Not required.

## Acknowledgements

The authors would like to express special thanks to Ms. Kirsi Ristkari for her assistance in the laboratory. The DDCB unit at the Faculty of Pharmacy (University of Helsinki), supported by HiLIFE and Biocenter Finland, is gratefully acknowledged for providing access to microbiology facilities and instruments. The Helsinki University Library supported the open-access publication of this article.

## Consent to Publish

Not applicable.

## Consent to Participate

Not applicable.

## Clinical Trial

Not applicable.

## Notes

### Competing Interest Statement

The authors have declared no competing interest.

